# Adaptation of eukaryotic membrane homeostasis to species-specific cellular lipid landscapes

**DOI:** 10.1101/2025.11.06.686945

**Authors:** Elisa Gomez-Gil, Paolo Guerra, Ying Gu, Sherman Foo, Benedicte Billard, Gabor Balogh, Maria Peter, Snezhana Oliferenko

## Abstract

The physicochemical properties of biological membranes must be maintained within a range compatible with cellular physiology. In the face of external perturbations, membrane lipid homeostasis mechanisms sense and control membrane features. How such mechanisms evolve to function in organisms with different cellular lipid make-up is unknown. Here, we address this fundamental question by exploiting the natural divergence in membrane lipid composition between the related fission yeasts *S. pombe* and *S. japonicus*. Using lipidomics and transcriptomics, we show that the activity of the membrane-bound transcriptional activator Mga2, which regulates the Δ-9 desaturase Ole1 expression, is set to sense distinct levels of membrane unsaturation in the two species. Through retro-engineering and physiological experiments, we identify an evolutionary divergent but functionally constrained element within the juxtamembrane region of Mga2, which fine-tunes its performance to species-specific lipid composition. Our experiments indicate that high baseline expression of *ole1*, set by cis-regulatory elements in its upstream non-coding region, has redefined the dynamic range of Mga2 activation in *S. pombe*, supporting high lipidome unsaturation. Our work explores an “experiment of nature” to highlight the broad principles underlying the organisation and evolution of membrane homeostasis, which should be applicable to other genetic networks supporting cellular homeostatic processes.

## Introduction

Biological membranes define cells and organelles, coordinate inter-organellar and cellular communication, and contribute greatly to cellular signalling and metabolism. Membrane physicochemical properties are shaped by both membrane lipid composition and membrane-associated proteome. These properties must be tightly regulated to meet the physiological demands of cells and organisms (Ernst et al. 2016; Harayama and Riezman 2018).

Eukaryotic lipid bilayers are based on glycerophospholipids of remarkable chemical diversity arising from different combinations of polar head groups and two fatty acyl (FA) chains. Variations in FA length and degree of unsaturation strongly influence membrane physicochemical properties. For example, cis double bonds cause kinks in the FAs, preventing tight lipid packing and hence, increasing membrane fluidity (Harayama and Riezman 2018). External cues such as variation in dietary lipid uptake or fluctuating temperature in poikilothermic organisms may greatly impact membrane properties, either through perturbing cellular lipid composition or directly affecting lipid diffusion (De Mendoza and Pilon 2019). To maintain cellular function under variable environmental conditions, organisms utilise homeostatic mechanisms that can sense membrane states and mount adaptive responses (Ernst et al. 2016). The key objective in this context is to maintain membrane physicochemical properties rather than the exact lipid composition.

Although the lipid metabolic toolbox may vary even in closely related species, organisms have evolved a set of conserved membrane sensing mechanisms to maintain membrane homeostasis. Those are usually based on transcriptional circuits (Ernst et al. 2016; Ernst et al. 2018; De Mendoza and Pilon 2019). A fascinating question is how homeostatic mechanisms evolve, given that they may need to sense and restore membranes to distinct optimal states in different organisms.

We address this question by exploiting two related fission yeast species, *Schizosaccharomyces pombe* and *Schizosaccharomyces japonicus*, which have profoundly different membrane glycerophospholipid composition. *S. pombe* membranes are massively unsaturated and relatively disordered, with a large proportion of glycerophospholipids where both FA chains are typically 18 carbon atoms long and mono-unsaturated at position Δ9. In contrast, *S. japonicus* synthesizes abundant asymmetrical saturated glycerophospholipids, containing a long FA (usually C18:0) at the *sn-1* position and a medium FA (C10:0) at the *sn-2* position of the glycerol backbone. *S. japonicus* also synthesizes unsaturated glycerophospholipids, albeit to a considerably lower extent than *S. pombe* (Makarova et al. 2020; Rao et al. 2025).

In fungi, FA unsaturation is controlled by the transcriptional activator Mga2 that regulates the expression of the essential Δ9-desaturase *ole1,* among other genes involved in lipid metabolism (Burr et al. 2016; Radanović et al. 2018; Marešová et al. 2024). Mga2 is produced as a precursor tethered to the endoplasmic reticulum (ER) membrane via the C-terminal transmembrane helix (TMH). The TMH senses membrane lipid saturation. When membranes become more saturated, Mga2 is ubiquitinylated and proteolytically cleaved via the proteasome (Hoppe et al. 2000). The liberated N-terminal transcriptionally active domain then drives the expression of *ole1* and other target genes in the nucleus, likely by modulating local chromatin accessibility (Burr et al. 2016; Ballweg and Ernst 2017).

Here, we show that the activation threshold and sensitivity of Mga2 have adjusted to the distinct ground states of membrane unsaturation in *S. pombe* and *S. japonicus* and identify a functionally constrained domain in the juxtamembrane region of Mga2 underlying these critical differences. Our data indicate that cis-regulatory elements in the untranslated region upstream of *S. pombe ole1* driving its high basal expression may have reshaped the dynamic range of Mga2 activation in this fission yeast. Together, our findings illustrate how conserved regulatory circuits can be rewired to meet species-specific physiological requirements, providing a broader framework for understanding the robustness and evolvability of other cellular homeostatic networks.

## Results

### The fatty acid desaturase *ole1* is a conserved target of Mga2-controlled transcriptional programmes in *S. pombe* and *S. japonicus*

To understand how the two species with drastically different membrane lipidomes adapt their glycerophospholipid composition to maintain membrane properties, we cultured *S. pombe* and *S. japonicus* cells across a range of physiological growth temperatures and performed shotgun electrospray ionization mass spectrometry (ESI-MS) analyses of total lipid extracts. Interestingly, we observed virtually orthogonal strategies of adaptation to growth temperature between the two fission yeasts (Fig. 1a and Supplemental Data 1, Tables 1, 2). While the *S. pombe* membrane glycerophospholipids remained highly unsaturated across the entire growth temperature range, *S. japonicus* reduced global glycerophospholipid unsaturation with increasing temperature. Conversely, whereas *S. japonicus* maintained high levels of glycerophospholipid asymmetry at all temperatures, *S. pombe* gradually increased the proportion of asymmetrical glycerophospholipids with decreasing temperature (Fig. 1b and Supplemental Data 1, Table 1). These results indicated that in changeable environment, cells favour the regulation of minor rather than major lipidome features. Additionally, this observation supports the hypothesis that asymmetrical saturated and symmetrical unsaturated glycerophospholipids confer similar physicochemical properties to lipid bilayers (Smith et al. 2021; Panconi et al. 2023; Rao et al. 2025).

**Figure 1.**
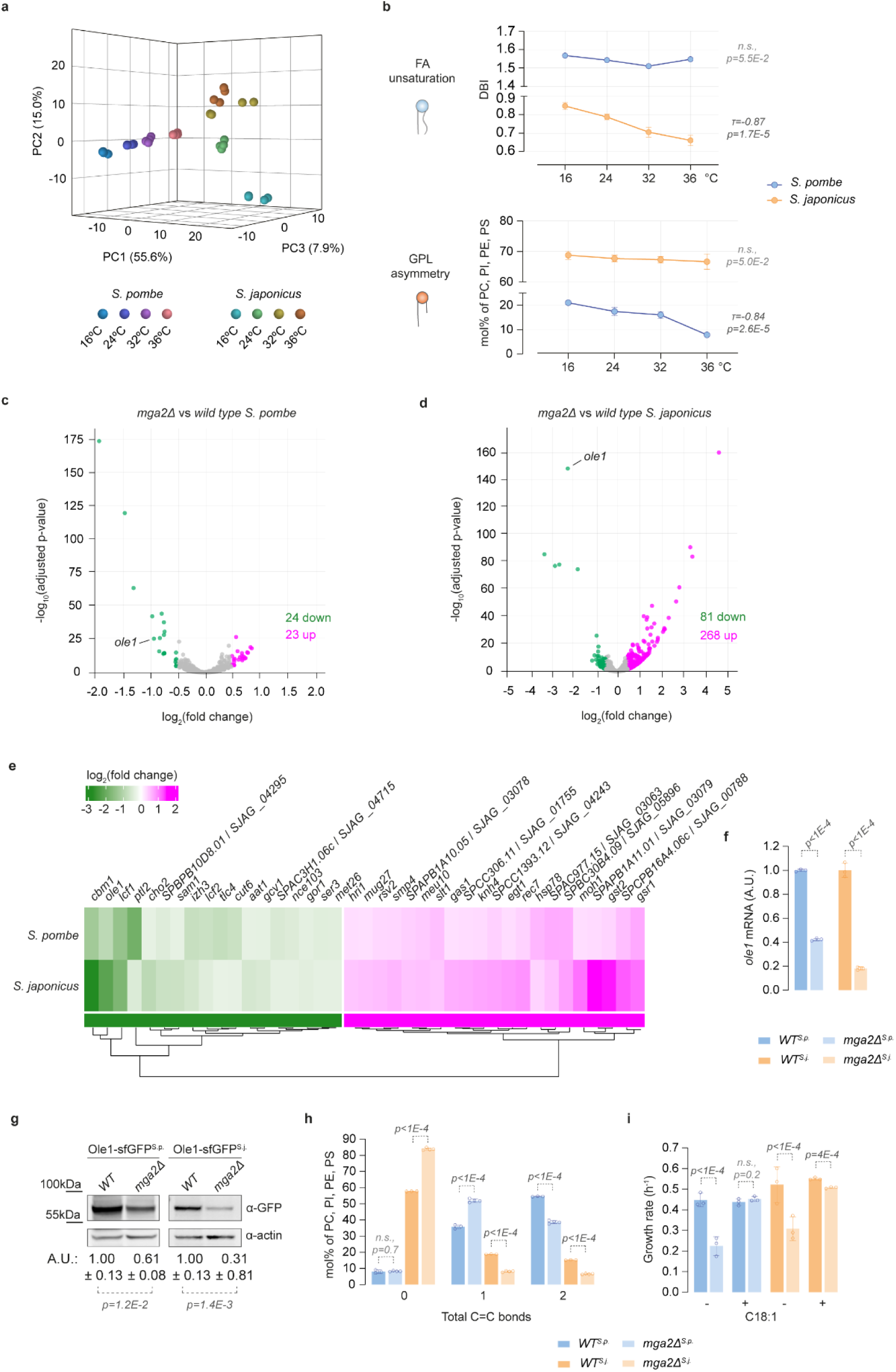
The fatty acid desaturase Ole1 is a conserved target of Mga2-mediated transcriptional regulation in *S. pombe* and *S. japonicus*. (**a**) Three-dimensional principal component analysis (PCA) plot of *S. pombe* and *S. japonicus* lipidomes grown in Edinburgh Minimal Media (EMM) at indicated temperatures. (**b**) Comparison of double bond indices (DBI) (top panel) and average combined fatty acid (FA) length (bottom panel) calculated for the sum of the four main glycerophospholipid (GPL) classes (phosphatidylcholine (PC), phosphatidylinositol (PI), phosphatidylethanolamine (PE) and phosphatidylserine (PS)) in *S. pombe* and *S. japonicus* wild-type cells grown in EMM at indicated temperatures. Data are represented as average ± S.D. Correlation coefficient (τ) and p-values are derived from Kendall’s rank correlation test (two-tailed). (**c**, **d**) Volcano plots showing RNAseq data from *S. pombe* (**c**) and *S. japonicus* (**d**) *mga2Δ* mutant vs wild type experiments. Genes significantly downregulated (green) or upregulated (magenta) are highlighted (-0.5 > log_2_(fold change) > 0.5 and adjusted p-value < 0.05). The position of *ole1* is indicated. (**e**) Heatmap showing a subset of differentially expressed genes commonly downregulated (green) or upregulated (magenta) in *mga2Δ S. pombe* and *S. japonicus* cells vs wild type. Colour bar showing log_2_(fold changes) is included. (**f**) Steady-state *ole1* mRNA levels in *S. pombe* and *S. japonicus* wild type (*WT*) and *mga2Δ* strains grown in YES, as measured by qPCR, normalized to the *WT*. (**g**) Western blot of Ole1-sfGFP in *S. pombe* and *S. japonicus WT* and *mga2Δ* strains grown in YES. Actin was used as loading control. Quantification of Ole1 protein levels is indicated under the blots, expressed as arbitrary units, normalized to the *WT*. (**h**) Grouping of GPL species according to the number of double bonds calculated for the sum of PC, PI, PE, and PS in *S. pombe* and *S. japonicus WT* and *mga2Δ* strains grown in EMM. Data are represented as average ± S.D. (n=2 biological, 1 technical repeats for *WT*; 2 biological, 2 technical repeats for *mga2Δ*). (**i**) Comparison of growth rates of *S. pombe* and *S. japonicus WT* and *mga2Δ* cells grown in YES in the absence or presence of 0.5 mM of the monounsaturated FA oleic acid (C18:1). (**a**, **b**) n=2 biological, 2 technical repeats. (**f**, **g**, **i**) Data are represented as average ± S.D. (n=3 biological repeats). (**f**-**i**) p-values are derived from two-tailed unpaired t-test.

Given that the two fission yeasts have drastically different fractions of unsaturated glycerophospholipids and distinct ways to regulate them in a temperature-dependent manner, we wondered whether the transcriptional programmes controlled by the Mga2 homeostatic pathway (Burr et al. 2016; Radanović et al. 2018; Marešová et al. 2024) are conserved. To address this question, we generated *S. pombe* and *S. japonicus* mutants lacking Mga2 and performed mRNA sequencing (RNA-seq) analysis. The number of differentially expressed genes in the absence of Mga2 was substantially higher in *S. japonicus* than in *S. pombe* (Fig. 1c, d, Supplemental Data 2, Tables 1, 2). The expression of the Δ9-desaturase *ole1* was significantly downregulated in *mga2Δ* cells in both fission yeasts, suggesting that *ole1* is a conserved target of this pathway (Fig. 1c, d). We detected additional differentially expressed genes, both down and upregulated, which overlapped between *S. pombe* and *S. japonicus mga2Δ* mutants, although the extent of their deregulation was different. The downregulated transcripts included previously described Mga2 targets in *S. pombe* related to lipid metabolism, including the acetyl-coA carboxylase *cut6*, the triacylglycerol lipase *ptl2* and the long-chain fatty acyl-coA synthetases *lcf1* and *lcf2* (Převorovský et al. 2015; Burr et al. 2017; Marešová et al. 2024). The commonly upregulated transcripts included two genes with roles in cell wall biogenesis (the essential 1,3-beta-glucanosyltransferase *gas1* and the regulator of 1,3-beta-glucan synthesis *gsr1*) (Fig. 1e).

Despite sharing a set of commonly regulated genes, *S. pombe* and *S. japonicus* lacking Mga2 also exhibited species-specific transcriptional responses, as evidenced by gene ontology (GO) enrichment analysis. For instance, in *S. pombe* the downregulated targets, which likely constitute the functional output of the Mga2 pathway, were associated with “molecular function” and “biological process” GO terms related to fatty acid metabolism. In *S. japonicus* such GO terms were associated with transmembrane transport processes (Fig. S1a and Supplemental Data 2, Tables 3, 4). In line with that, “cellular component” GO terms in *S. pombe* included the “endoplasmic reticulum” and “lipid droplet”, but in *S. japonicus* the only enriched GO term was “plasma membrane” (Fig. S1a and Supplemental Data 2, Tables 3, 4). The upregulated transcripts in *S. pombe* were enriched for unfolded protein responses, whereas in *S. japonicus* the strongest enrichments were associated with hydrolase activity, cell wall biogenesis and phospholipid catabolic process (Fig. S1b and Supplemental Data 2, Tables 3, 4).

Recent studies in *S. pombe* suggest that Mga2 cooperates with the CBF1/Su(H)/Lag-1 (CSL) family transcription factor Cbf11 to promote the expression of lipid metabolism genes, with Cbf11 binding the DNA and Mga2 acting as a transcriptional activator (Marešová et al. 2024). Consistent with this model, we detected significant enrichment of the Cbf11 binding motif (Převorovský et al. 2015) in the promoters of downregulated targets in *S. pombe* (Supplemental Data 2, Table 5). Such enrichment was not observed in *S. japonicus* (Supplemental Data 2, Table 6). Nevertheless, we identified putative Cbf11 binding sites within the 1 kb upstream regions of *ole1* and other Mga2 lipid metabolism-related targets in both sister species. In *S. japonicus* the putative regulatory network expanded to genes involved in other cellular processes, including DNA replication, recombination and repair, cell-cell conjugation and fusion (Supplemental Data 2, Tables 5, 6).

Our results so far indicated that despite the divergence in the transcriptional output of the Mga2 pathway and/or cellular consequences associated with its deficiency, a subset of Mga2 targets involved in lipid metabolism was conserved between the two species. Related to FA unsaturation, reverse-transcription qPCR (RT-qPCR) analysis confirmed that Mga2 regulated *ole1* expression in both species, albeit to different extent. While *ole1* expression in *S. japonicus* depended critically on Mga2, its contribution to *ole1* regulation was less pronounced in *S. pombe* (Fig. 1f). Notably, *ole1* transcript levels correlated well to the protein abundance in both species, as shown by Western blotting and microscopy of Ole1 tagged C-terminally with superfolder GFP (sfGFP) (Fig. 1g and Fig. S1c).

Consistent with the interspecies difference in *ole1* regulation, comparative ESI-MS lipidomics revealed a greater impact of Mga2 deficiency in *S. japonicus*, with most glycerophospholipids becoming fully saturated and asymmetrical, while *S. pombe* responded by producing more mono-unsaturated vs di-unsaturated glycerophospholipid species (Fig. 1h, Fig.S1d, g, h and Supplemental Data 1, Table 3). The lack of Mga2 caused further alterations to the cellular lipid landscape. We observed an increased ratio between phosphatidylethanolamine (PE) and the sum of phosphatidylcholine and phosphatidylinositol (PE/(PC + PI)) in *mga2Δ* cells as compared to the wild type in both species (Fig. S1e, f). Unlike PC and PI that have a cylindrical shape and form flat lipid bilayers, PE is conically shaped, generating intrinsic negative curvature. Thus, a relative increase in a non-bilayer PE may indicate the need to maintain the collective physical properties of the membrane when FA unsaturation is critically reduced (Renne and De Kroon 2018; Yang et al. 2025). Besides other adjustments in minor lipid species, *S. japonicus mga2Δ* cells showed a significant reduction in the levels of diacylglycerol (DG) and triacylglycerol (TG) (Fig. S1i). The latter could be due to the upregulation of genes involved in lipid catabolic processes (Fig. S1b). On the other hand, *S. pombe* cells lacking Mga2 exhibited increased levels of phosphatidic acid (PA) and inositol phosphoceramide (IPC) (Fig. S1i).

The lack of Mga2 strongly impaired the growth rate of both fission yeasts. Suggesting that reduction in glycerophospholipid unsaturation could be at the root of this phenotype, supplementation of *mga2Δ* cells with exogenous unsaturated FA, oleic acid (C18:1), restored their growth rate to wild-type levels in *S. pombe*, and led to almost complete rescue in *S. japonicus* (Fig. 1i, also see (Burr et al. 2016) for *S. pombe*). Thus, *ole1* is a critical target of Mga2 in both species.

Together, these findings indicate that despite the evolutionary rewiring of Mga2-dependent transcriptional response, the Mga2-Ole1 relationship is conserved between *S. pombe* and *S. japonicus*. However, Ole1 is under a stronger Mga2 regulation in *S. japonicus*, suggesting that this regulatory circuit may have adapted to the distinct levels of glycerophospholipid unsaturation in these organisms.

### Mga2 activity is tuned to sense different membrane states in the two species

To investigate how Mga2 interprets distinct membrane lipid environments and differentially regulates *ole1* expression, we examined its activation by proteolytic cleavage in *S. pombe* and *S. japonicus* (Hoppe et al. 2000; Burr et al. 2017). To this end, we N-terminally tagged Mga2 at its endogenous locus with GFP (GFP-Mga2), allowing us to detect both the ER membrane-bound precursor (P) and the transcriptionally active N-terminal fragment (N). Strikingly, whereas at steady state *S. japonicus* showed a considerable fraction of uncleaved precursor in the ER, Mga2 in *S. pombe* was largely cleaved, with its N-terminal fragment localizing to the nucleus (Fig. 2a, b). Treatment with the proteasome inhibitor bortezomib resulted in strong accumulation of the Mga2 precursor in both fission yeasts, suggesting that the proteasome-dependent proteolytic processing of Mga2 was conserved (Fig. S2a, b, also see (Burr et al. 2017)). Constitutive Mga2 cleavage may sustain extremely high levels of lipidome unsaturation required by *S. pombe* (Fig. 1b, i, Fig. S1d).

**Figure 2.**
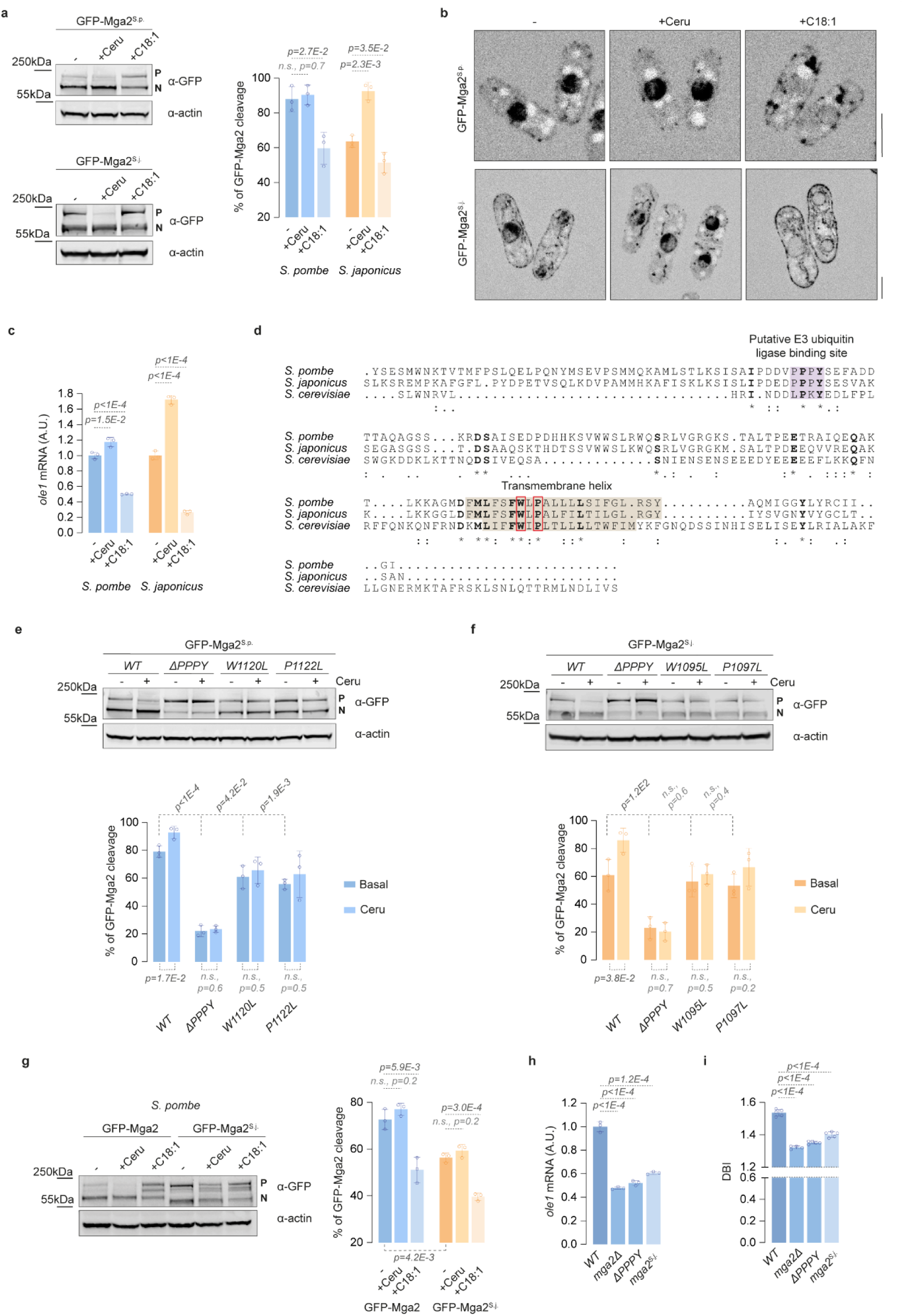
*S. pombe* and *S. japonicus* Mga2 proteins exhibit distinct activation thresholds and sensitivities. (**a**) Western blot of *S. pombe* (top panel) and *S. japonicus* (bottom panel) cells expressing GFP-Mga2 grown in YES in the absence or the presence of 10 µM cerulenin for 1 hour (Ceru) or 0.5 mM of the unsaturated FA oleic acid for 1 hour (C18:1). Actin was used as a loading control. The right panel shows the quantification of GFP-Mga2 cleavage. (**b**) Single plane spinning disk confocal images of *S. pombe* and *S. japonicus* cells from experiment shown in (**a**). Scale bars (5 µm) for each species are shown on the right. (**c**) *ole1* mRNA levels in *S. pombe* and *S. japonicus* wild type cells grown in YES in the absence or the presence of cerulenin or oleic acid as described in (**a**), as measured by qPCR, normalized to the steady-state conditions for each species. (**d**) ClustalW alignment of the C-terminal region (last 160 amino acids) of Mga2 proteins in *S. pombe*, *S. japonicus* and *S. cerevisiae* represented using ESPript 3.0. ‘*’: identical amino acid, ‘:’: conserved substitution, ‘.’: semi-conserved substitution. The putative conserved residues involved in E3 ligase binding are shaded in purple. The predicted transmembrane helices are shaded in brown. The conserved residues involved in sensing membrane lipid packing in *S. cerevisiae* are boxed in red. (**e, f**) Western blots of *S. pombe* (**e**) and *S. japonicus* (**f**) cells expressing the wild type or indicated mutant versions of GFP-Mga2 grown in YES in the absence or the presence of 10 µM cerulenin for 1 hour (Ceru). Actin was used as a loading control. The bottom panels show the quantification of GFP-Mga2 cleavage. (**g**) Western blot of *S. pombe* cells expressing the wild type or the *S. japonicus* version of GFP-Mga2 grown in YES in the absence or the presence of 10 µM cerulenin (Ceru) for 1 hour or 0.5 mM of oleic acid for 1 hour (C18:1). Actin was used as a loading control. The right panel shows the quantification of GFP-Mga2 cleavage. (**h**) Steady-state *ole1* mRNA levels in *S. pombe* strains of the indicated genotypes grown in YES, as measured by qPCR, normalized to the wild type (*WT*). (**i**) Comparison of DBI calculated for the sum of the four main GPL classes (PC, PI, PE and PS) in *S. pombe* strains of the indicated genotypes grown in EMM. Data are represented as average ± S.D. (n=3 biological, 2 technical repeats for *WT*, *ΔPPPY* and *mga2^S.j.^*; 2 biological, 2 technical repeats for *mga2Δ*). (**a**, **e*-*g**) P and N indicate precursor and cleaved N-terminal transcriptional activator forms, respectively. Mga2 cleavage is calculated as N / (N + P), expressed as percentage. (**a**, **c**, **e**-**h**) Data are represented as average ± S.D. (n=3 biological repeats). (**a**, **c**, **e**-**i**) p-values are derived from two-tailed unpaired t-test.

The distinct steady-state activation patterns of Mga2 prompted us to examine whether the sensitivity of Mga2 to alterations in fatty acid unsaturation also differs between *S. pombe* and *S. japonicus*. The two yeasts responded differently to the treatment with cerulenin, which inhibits de novo FA synthesis and thus prevents FA desaturation, likely increasing lipid packing (Awaya et al. 1975; Omura 1976; Burr et al. 2017). Whereas cerulenin treatment induced GFP-Mga2 proteolytic cleavage and nuclear translocation in *S. japonicus*, it had a negligible impact on GFP-Mga2 processing in *S. pombe* - in line with most GFP-Mga2 being cleaved already under normal conditions (Fig. 2a, b). Conversely, supplementation with unsaturated oleic acid inhibited GFP-Mga2 cleavage in both yeasts (Fig. 2a, b). GFP-Mga2 cleavage largely correlated with *ole1* expression levels as measured by RT-qPCR (Fig. 2c), although *ole1* expression appears to be a more sensitive readout of Mga2 pathway activation as compared to measuring Mga2 cleavage. Importantly, both decreasing and increasing unsaturated fatty acids elicited a stronger regulation of *ole1* expression in *S. japonicus* than in *S. pombe* (Fig. 2c). We concluded that Mga2 proteins in the two species exhibit distinct activation thresholds and different degree of sensitivity.

We next wondered whether the distinct activation patterns of *S. pombe* and *S. japonicus* Mga2 proteins could be attributed to differences in their lipid saturation-sensing mechanism (Covino et al. 2016). Binding of the E3 ubiquitin ligase Rsp5 was shown to be critical for proteasome-dependent Mga2 processing in budding yeast. This site and its relative position to the transmembrane helix are conserved in both fission yeasts (Fig. 2d). The deletion of this putative E3 ubiquitin ligase binding site (ΔPPPY) largely abolished Mga2 cleavage both under normal conditions and upon cerulenin treatment in both *S. pombe* and *S. japonicus* (Fig. 2e, f and Fig. S2c, d).

The sequences of Mga2 transmembrane helix are also well-conserved between both fission yeasts and the evolutionarily distant *S. cerevisiae*, including the two residues (W1042 and P1044) shown to sense membrane saturation and enable efficient proteolytic activation in budding yeast (Covino et al. 2016) (Fig. 2d). Mutation of either of the corresponding residues in *S. pombe* (W1120, P1122) and *S. japonicus* (W1095, P1097) to Leucine, significantly reduced Mga2 cleavage as compared to the wild type, and abolished its induction in response to cerulenin treatment (Fig. 2e, f and Fig. S2c, d). Thus, it appears that the mechanism by which Mga2 transmembrane helix senses lipid unsaturation is broadly conserved and therefore cannot explain the different activation thresholds between the two sister species.

To probe if other regions of the Mga2 sequence are responsible for its distinct cleavage patterns, we replaced the endogenous *mga2* ORF in *S. pombe* with the *S. japonicus* ortholog at the native locus. The swapped protein was N-terminally tagged with GFP (GFP-Mga2^S.j.^). Intriguingly, *gfp-mga2^S.j^*mutant *S. pombe* showed markedly reduced Mga2 cleavage at steady-state as compared to the endogenous version (Fig. 2g and Fig. S2e). Reduced Mga2 cleavage correlated with decreased *ole1* expression and a concomitant reduction in global glycerophospholipid unsaturation, similar to the *mga2Δ* mutant or the mutant lacking the putative E3 ubiquitin ligase binding site (Fig. 2h, i). The transplanted *S. japonicus* Mga2 responded only to the decrease in membrane order upon supplementation with oleic acid, but not to cerulenin treatment (Fig. 2g and Fig. S2e).

Our results indicate that the *S. japonicus* Mga2 that has evolved in an organism with relatively low membrane lipid desaturation, is “satisfied” in *S. pombe* membrane environment. In this species, even in the absence of endogenous Mga2 or upon cerulenin treatment, the lipidome unsaturation remains high enough (Fig. S1d), and thus, lies far above the normal *S. japonicus* Mga2 sensitivity range. On the other hand, the system remains sensitive to extreme desaturation triggered by supplementation with oleic acid. We concluded that the activation threshold is determined by the Mga2 protein sequence itself.

### Differences in the juxtamembrane region of Mga2 are responsible for its species-specific activation patterns

Since Mga2 functions homeostatically to maintain membrane properties, the divergence in the transcriptional output of the Mga2 pathway could contribute to its distinct activation patterns. To explore this possibility, we generated chimeric Mga2 proteins in *S. pombe* and *S. japonicus* by swapping the first 130 amino acids corresponding to their N-terminal transcriptional activator domains (Hoppe et al. 2000; Romanauska and Köhler 2021) in the native chromosomal context (Mga2-TAswap, Fig. 3a), and analysed changes in global mRNA abundance by RNA-seq. We did not observe any differentially expressed genes in *S. japonicus* mutant as compared to the wild type. The *S. pombe* Mga2-TAswap^S.j.^ mutant showed only two mildly downregulated genes, the acid phosphatase *pho1* and the adenine-repressible gene *rds1*, neither related to lipid metabolism. In addition, we observed an increase in the transcripts of *mga2* itself and the adjacent gene *pot1*, likely as a consequence of the engineering at this locus (Fig. 3b). Of note, both *S. pombe* and *S. japonicus* Mga2-TAswap mutants maintained nearly wild type-like *ole1* expression levels and, unlike the *mga2Δ* mutants, the chimeric strains did not display growth defects (Fig. S3a, b). These results indicate that the transcriptional activator domains of *S. pombe* and *S. japonicus* Mga2 proteins can substitute for each other, suggesting that the specificity of Mga2 activation patterns must reside somewhere else within the protein.

**Figure 3.**
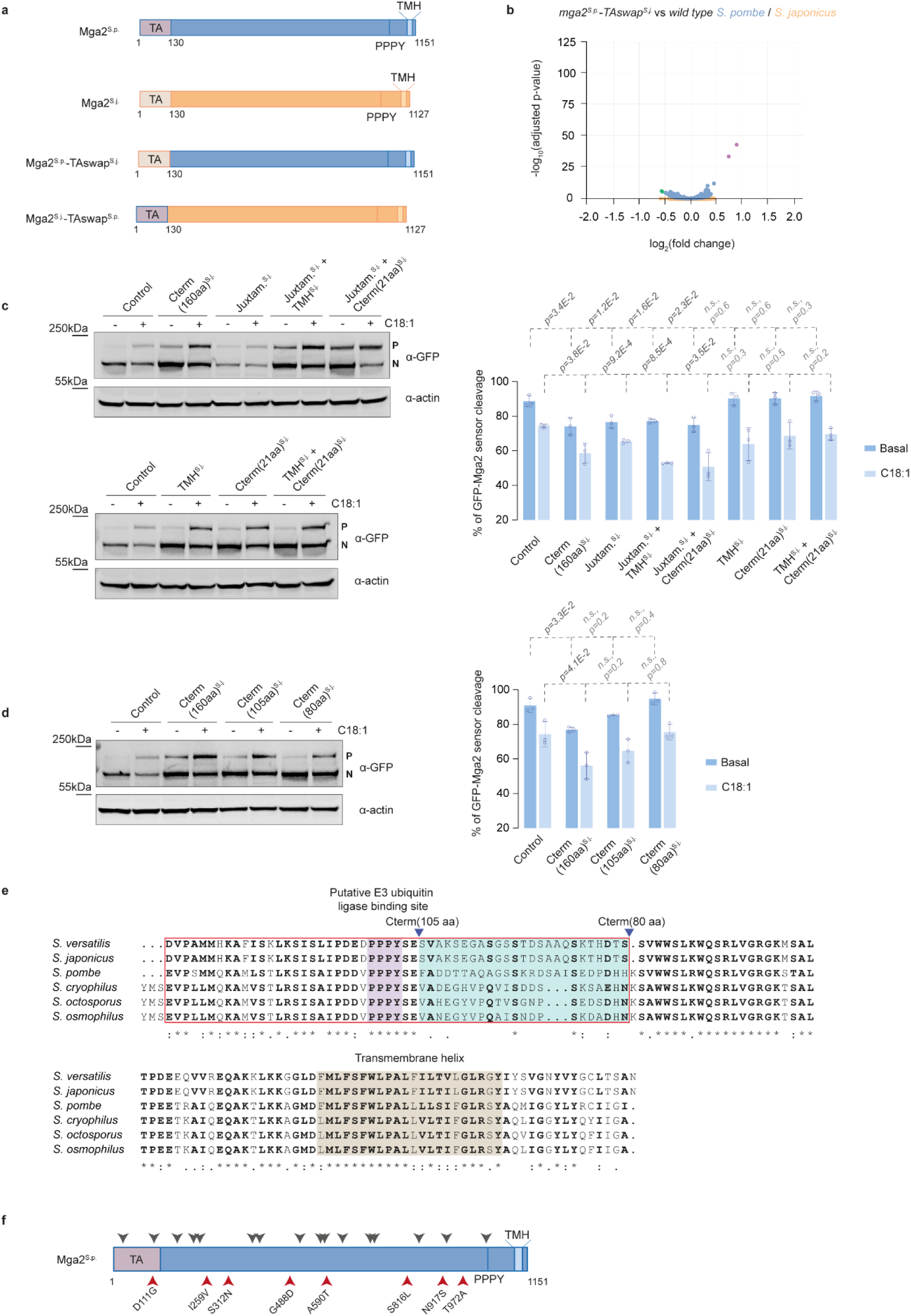
Mga2 species-specific activation pattern is determined by its juxtamembrane domain. (**a**) Diagram of *S. pombe* and *S. japonicus* Mga2 proteins and chimeric constructs. Mga2^S.p.^ and Mga2^S.j.^: wild type Mga2 protein showing the transcriptional activator domain (TA), the putative E3 ligase binding site (PPPY) and the predicted transmembrane helix (TMH) for *S. pombe* and *S. japonicus*, respectively. Mga2^S.p.^-TAswap^S.j.^: *S. pombe* Mga2 version with the *S. japonicus* transcriptional activator domain. Mga2^S.j.^-TAswap^S.p.^: *S. japonicus* Mga2 version with the *S. pombe* transcriptional activator domain. (**b**) Volcano plot showing RNA-seq data from *S. pombe* (blue) and *S. japonicus* (orange) Mga2-TAswap mutant vs wild type experiments. *S. pombe* genes significantly downregulated (green) or upregulated (magenta) are highlighted (-0.5 > log_2_(fold change) > 0.5 and adjusted p-value < 0.05). Note that there were no significantly regulated genes in *S. japonicus*. (**c**) Western blots of *S. pombe* cells expressing GFP-Mga2 sensor constructs with the indicated chimeric C-terminal domains grown in YES in the absence or the presence of 1 mM oleic acid (C18:1) for 3 hours. Actin was used as a loading control. The right panel shows the quantification of GFP-Mga2 cleavage. (**d**) Western blot of *S. pombe* cells expressing GFP-Mga2 sensor constructs with the indicated chimeric C-terminal domains grown in YES in the absence or the presence of 1 mM oleic acid (C18:1) for 3 hours. Actin was used as a loading control. The right panel shows the quantification of GFP-Mga2 cleavage. (**e**) ClustalW alignment of the C-terminal region (last 135 amino acids) of Mga2 proteins in the six species of the *Schizosaccharomyces* genus: *S. versatilis*, *S. japonicus*, *S. pombe*, *S. cryophilus*, *S. octosporus* and *S. osmophilus* represented using ESPript 3.0. ‘*’: identical amino acid, ‘:’: conserved substitution, ‘.’: semi-conserved substitution. The putative conserved residues involved in E3 ubiquitin ligase binding are shaded in purple. The most divergent stretch of residues across species, corresponding to C terminal positions 105 to 80, is shaded in cyan. The predicted transmembrane helix domains are shaded in brown. The region responsible for Mga2 species-specific activation patterns is boxed in red. (**f**) Diagram of *S. pombe* Mga2 showing the synonymous (grey) and non-synonymous (red) variations found in 57 natural isolates of diverse origins. The transcriptional activator domain (TA), the putative E3 ligase binding site (PPPY) and the predicted transmembrane helix (TMH) are indicated. (**c**, **d**) P and N indicate precursor and cleaved N-terminal transcriptionally active forms, respectively. Mga2 cleavage is calculated as N / (N + P), expressed as percentage. Data are represented as average ± S.D. (n=3 biological repeats). p-values are derived from two-tailed unpaired t-test.

We next explored the role of the entire C-terminal “sensor” domain of Mga2, which includes the last 160 amino acids of the protein and comprises the juxtamembrane domain, the transmembrane helix (TMH), and the extreme C-terminus that is predicted to form an amphipathic helix (Fig. 2d). The juxtamembrane domain contains the putative E3 ubiquitin ligase binding site followed by an intrinsically disordered region that extends to the TMH (Ballweg et al. 2020). We engineered *S. pombe* Mga2-based constructs inspired by the LipSat sensors developed in budding yeast (Romanauska and Köhler 2021). These N-terminally GFP-tagged sensors lacked the 130 amino acid-long transcriptional activator domains, uncoupling membrane order sensing from the expression of *ole1* and other Mga2 targets. To avoid hetero-dimerization with the endogenous Mga2 protein (Romanauska and Köhler 2021), these constructs were integrated at the *mga2* locus and were the only source of Mga2 in the mutant cells. Consistent with reduced lipidome unsaturation in the absence of Mga2 transcriptional activity, the control *S. pombe* Mga2 sensor was predominantly cleaved and localized to the nucleus at steady state (Fig. 3c, Fig. S3c). In line with its ability to report on fatty acyl chain unsaturation in the ER membrane, sensor processing was strongly reduced upon exogenous supplementation with oleic acid, leading to its increased retention in the ER (Fig. 3c, Fig. S3c). Interestingly, the sensor construct, in which the *S. pombe* C-terminal domain was replaced by the corresponding 160 amino acids from *S. japonicus* (Cterm(160aa) ^S.j.^) showed reduced cleavage even at steady state, which was further decreased upon supplementation with oleic acid (Fig. 3c, Fig. S3c). This behaviour closely resembled the *S. japonicus* Mga2 activation pattern, suggesting that the features responsible for species-specific cleavage are encoded within the C-terminal 160 amino acids of Mga2.

To narrow down the sequence responsible for this phenotype, we generated additional *S. pombe* sensor constructs carrying the *S. japonicus* juxtamembrane (Juxtamembrane^S.j.^), TMH (TMH^S.j.^) or C-terminal amphipathic helix (Cterm(21aa) ^S.j.^), either individually or in combination, and tested whether these chimeras sensed membrane environment in a *S. pombe*-like or *S. japonicus*-like fashion. Strikingly, all sensor constructs that included the juxtamembrane region, either alone or combined with the TMH or the amphipathic helix, showed *S. japonicus*-like activation pattern, making the sensor less prone to cleavage than the control construct already at steady state. In contrast, the chimeras containing only the TMH, the C-terminal amphipathic helix, or a combination of both displayed *S. pombe*-like phenotypes (Fig. 3c, Fig. S3c). We observed variability in protein abundance among the different constructs, likely due to differences in protein stability and/or half-life caused by protein engineering.

Following the same strategy and shortening the *S. japonicus* juxtamembrane domain sequence in our chimeric constructs, we successfully mapped a 55 amino acid long region responsible for the species-specific activation pattern (Fig. 3d, see boxed sequence in Fig. 3e, Fig. S3c). This segment includes the conserved putative E3 ubiquitin ligase binding site and a part of the intrinsically disordered region. Interestingly, the latter is the most evolutionarily divergent stretch of amino acids within the C-termini of *Schizosaccharomyces* Mga2 proteins. Yet, it is highly conserved between 57 *S. pombe* natural isolates of diverse origins (Jeffares et al. 2015) (Fig. 3e, f and Supplemental Data 3, Table 1), suggesting that it remains functionally constrained within an individual species. The chimeric mutant (Cterm(105aa) ^S.j.^) that lacked the E3 ubiquitin ligase binding site but contained this species-specific intrinsically disordered sequence also showed enhanced localization to the ER at steady state and further ER retention upon oleic acid treatment, although the differences in cleavage as compared to the control did not reach statistical significance (Fig. 3d, e and Fig. S3c).

We concluded that the 55 amino acid region within the juxtamembrane domain has evolved to function in different species-specific lipid environments, with *S. pombe* Mga2 operating in a highly unsaturated lipidome.

### High basal expression of *ole1* in *S. pombe* is facilitated by a distinct architecture of its upstream non-coding region

In *S. pombe*, even in the absence of Mga2, there is sufficient amount of Ole1 to drive higher lipidome unsaturation as compared to *S. japonicus* (Fig. 1 and Fig. S1). Consistently, the C-terminally tagged Ole1-sfGFP was much more abundant (∼5 times at the protein level and ∼2.5 times at the transcript level) in *S. pombe* than in its sister species (Fig. 4a, b, Fig. S4a).

**Figure 4.**
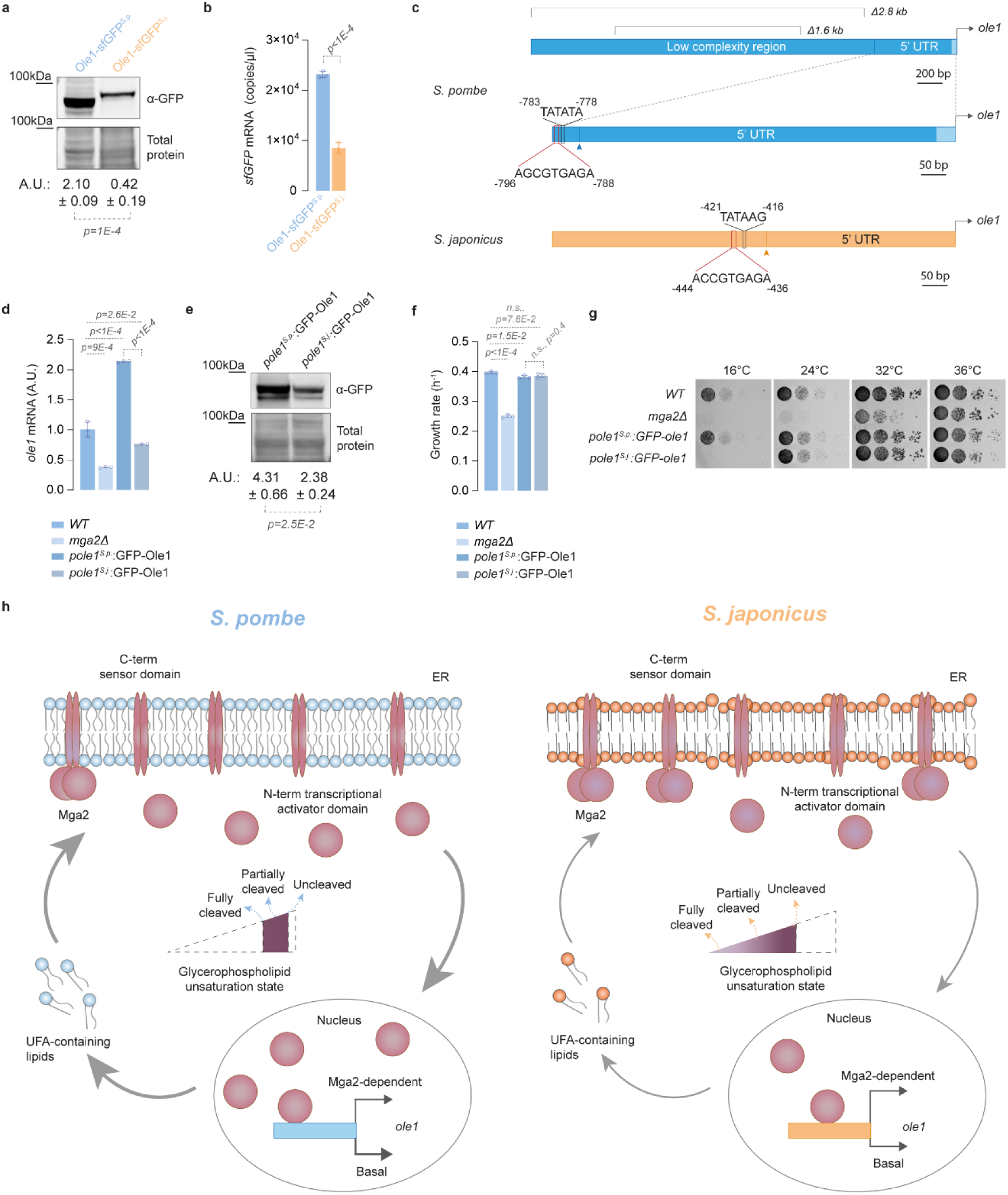
Mga2 functions at the upper limit of its dynamic range due to high baseline expression of Ole1 in *S. pombe*. (**a**) Western blot of *S. pombe* and *S. japonicus* cells expressing Ole1-sfGFP grown in YES. Revert 700 stain was used to visualize total protein loading. The quantification, expressed as arbitrary units, is shown below the blot. (**b**) Absolute quantification of Ole1-sfGFP mRNA levels in *S. pombe* and *S. japonicus* cells grown in YES, expressed as copies per µl. (**c**) Schematics of the non-coding region upstream of *ole1* ORF in *S. pombe* and *S. japonicus*. Putative Cbf11 binding sites are boxed in red. Putative TATA boxes are boxed in black. The location of transcription start sites (TSS) are indicated with arrows. (**d**) Steady-state *ole1* mRNA levels in *S. pombe* wild type (*WT*), *mga2Δ*, *pole1^S.p.^*:GFP-Ole1 and *pole1^S.j.^*:GFP-Ole1 strains grown in YES, as measured by qPCR, normalized to the *WT*. (**e**) Western blot of *pole1^S.p.^*:GFP-Ole1 and *pole1^S.j.^*:GFP-Ole1 *S. pombe* cells, grown in YES. Revert 700 stain was used to visualize total protein loading. The quantification, expressed as arbitrary units, is shown below the blot. (**f**) Comparison of growth rates of *S. pombe* strains of the indicated genotypes grown in YES at 30°C. (**g**) Serial dilution assay of *S. pombe* strains of the indicated genotypes carried out in YES medium at the indicated temperatures. (**h**) A pictorial model suggesting how the high levels of Mga2-independent baseline *ole1* expression and thus, high levels of lipidome desaturation, may force the *S. pombe* Mga2 to operate at the upper limit of its dynamic range. (**a**, **b**, **d**-**f**) Data are represented as average ± S.D. (n=3 biological repeats). p-values are derived from two-tailed unpaired t-test.

Divergence in cis-regulatory elements such as transcription factor binding sites, other intrinsic DNA sequence properties or distinct chromatin architecture may determine quantitative changes in gene expression that occur in evolution (Dalal et al. 2016; Kim and Wysocka 2023). In *S. pombe* the transcription start site (TSS) for *ole1* has been mapped to -747 base pairs (bp) upstream of ORF ((Li et al. 2015), consistent with our RNA-seq data). It is preceded by a canonical TATA box motif (TATATA) starting at the position -783 bp. A Cbf11-Mga2 binding site is located 14 bp upstream of the TATA box at the position -797 bp ((Marešová et al. 2024) and Fig. 4c, Supplemental Data 2, Table 5). In the absence of Mga2, the remaining *ole1* transcription was still initiated at the same TSS, suggesting high Mga2-independent baseline promoter activity (Fig. S4b).

Interestingly, virtually the entire non-coding sequence upstream of *ole1* ORF lies within a larger ∼3.4 kb low-complexity region rich in poly(dA:dT) tracts, which are absent at the equivalent positions in *S. japonicus* and other fission yeasts (Fig. 4c and Fig. S4c). The AT-rich sequences can be associated with nucleosome-depleted areas that increase the accessibility of regulatory promoter elements (Raveh-Sadka et al. 2012), although it appears that in *S. pombe*, the nucleosome-positioning factors may override biophysical DNA sequence properties (Lantermann et al. 2010). We wondered whether the presence of the low complexity region upstream of the putative promoter was required to maintain high basal *ole1* expression in *S. pombe*. Gene disruption experiments using *S. pombe* heterozygous diploids revealed that a 1.6 kb fragment from the central part of the low complexity region could be deleted without affecting *ole1* expression levels (Fig. S4d), whereas a longer 2.8 kb fragment comprising most of the region, including the putative *ole1* promoter containing the Cbf11-Mga2 binding site and the TATA-box was essential for cell viability (0 knock out mutants out of 32 germinated spores were recovered; see Methods, Fig. 4c). We concluded that the high levels of *ole1* expression required for viability in *S. pombe* are driven by the region between -1177 and -747 bp upstream of *ole1* ORF, which contains its promoter at positions -797 (Cbf11-Mga2 binding site) to -783 bp (TATA-box). Importantly, these results also suggested that there were no alternative promoters downstream of that unit, which could support *ole1* expression.

In *S. japonicus*, the non-coding region upstream of *ole1* ORF includes a similar set of *cis* elements although it is positioned closer to the start codon. The TSS, approximately at position -375 bp, is preceded by a relaxed TATA-box (TATAAG) at position -421 bp and a putative Cbf11-Mga2 binding site at position -444 bp (Fig. 4c, Supplemental Data 2, Table 6). Of note, whereas the 800 bp region upstream of the *S. pombe ole1* ORF, including its promoter, exhibited very high AT content (68%), the corresponding region in *S. japonicus* had only 55% AT.

To test if the quantitative differences in *ole1* expression between *S. pombe* and *S. japonicus* were due to the architecture of the upstream non-coding sequences, we have replaced the 800 bp of *S. pombe ole1* upstream of the ORF with the cognate *S. japonicus* sequence. We generated two *S. pombe* strains, in which the N-terminally GFP-tagged Ole1 was expressed either from the native 800 bp-long (*pole1^S.p^*.) regulatory region or the corresponding *S. japonicus* sequence (*pole1^S.j^*.). Both constructs were integrated at the native *ole1* locus. We noticed that the *ole1* transcript levels were approximately twice as high in the control *pole1^S.p^*.:GFP-Ole1 strain as compared to the wild type, likely due to differences in mRNA stability or translation because of tagging or locus engineering (Fig. 4d). In the same setup, using *pole1^S.j^*. resulted in a considerably lower *ole1* mRNA expression (∼2.5x lower; Fig. 4d), which translated to lower levels of GFP-Ole1 protein (Fig. 4e and Fig. S4e). The expression of *ole1* mRNA in *pole1^S^ ^j^*. cells reached intermediate levels between *S. pombe* wild type and the mutant lacking Mga2 (Fig. 4d). In line with that, whereas the *pole1^S.j^*.cells could grow normally within 24°C to 36°C temperature range, they failed to form colonies at 16°C, similar to the *mga2Δ* mutant (Fig. 4f, g).

Importantly, unlike in the wild type (Fig. 1) or in the *pole1^S. p^*. mutant, Mga2 has become essential in the *pole1^S. j^*. genetic background (0 double mutants out of 44 germinated spores were recovered; see Methods). These results suggested that the baseline activity of the *S. japonicus ole1* promoter could not support sufficient *ole1* expression to maintain the required levels of lipidome unsaturation in *S. pombe*. We concluded that the high AT-rich content or other sequence features within the upstream 800bp of *S. pombe ole1* allow for high basal expression of this gene.

Based on our findings, we favour a model where high basal, Mga2-independent, expression of *ole1* in *S. pombe* due to the evolutionary invasion of a low-complexity region in its upstream non-coding sequence has redefined the dynamic range of lipidome unsaturation sensed by Mga2. When the lipidome is highly unsaturated at steady state, the Mga2-dependent homeostasis mechanism can only operate at the upper limit of the parameter space, preventing extreme desaturation. We hypothesize that *S. pombe* has become “addicted” to massive glycerophospholipid desaturation, with high Mga2 basal cleavage/activation required to support it, leaving little capacity for further response. Conversely, in *S. japonicus* the expression of *ole1* is almost fully dependent on Mga2. In this case, Mga2 operates within a broader dynamic range, retaining a capacity for bi-directional regulation upon perturbations to membrane order. Thus, we hypothesize that the species-specific differences in the sequence of the Mga2 juxtamembrane domain determining its cleavage efficiency may have evolved to sustain distinct dynamic ranges of lipidome unsaturation (Fig. 4h).

## Discussion

Both in *S. pombe* and *S. japonicus*, Mga2 positively regulates the expression of a conserved set of genes related to lipid metabolism (Fig. 1c-e and (Burr et al. 2016; Marešová et al. 2024)). The *S. japonicus* Mga2 may have additional target genes, consistent with its broader impact on cellular physiology, as compared to *S. pombe* (Fig. 1c, d, i, and Fig. S1a, b). The rewiring of the Mga2 regulon could be due to the evolutionary reshuffling of the Cbf11 binding motifs, additional interactions of Mga2 with another transcription factor, or differences in other cis-regulatory elements or chromatin architecture at target genes (Li and Johnson 2010). Given that the transcriptional activator domains of *S. pombe* and *S. japonicus* Mga2 proteins can functionally substitute for each other (Fig. 3a, b and Fig. S3a, b), our data suggest that the trans-specificity of Mga2 is conserved whereas the cis-regulation may have been rewired. Also, the extent of the transcriptional response may be increased if new cis-elements arise in the regulatory sequences of genes encoding other transcriptional regulators. In line with that, only in *S. japonicus* but not *S. pombe*, the transcription factor TFIIH *tfb4* has a putative Cbf11-binding site and is positively regulated by Mga2 (Supplemental Data 2, Tables 5, 6).

Of note, the relationship between Mga2 and Cbf11 resembles that between the Notch receptors and the CSL transcription factors governing cell-cell communication and developmental decisions in metazoans (Bray 2006). Although the Notch receptor resides at the plasma membrane whereas the Mga2 precursor is located at the ER, both are single-pass transmembrane proteins activated through proteolytical cleavage, albeit via different mechanisms. In both cases, cleavage releases an ankyrin repeat containing fragment that translocates to the nucleus, where it interacts with a CSL-family transcription factor to regulate gene expression (Kovall 2007). Thus, similar regulatory logics may be harnessed in evolution to regulate different physiological processes.

The gene encoding the Δ9-desaturase *ole1* is a critical target of Mga2 in both species (Fig. 1 and Fig. S1). However, the operational settings for Mga2-dependent lipidome homeostasis in *S. pombe* and *S. japonicus* are different. Not only the two species differ profoundly in the extent of membrane unsaturation, but the *S. japonicus* membranes are also rich in asymmetrical glycerophospholipids (Makarova et al. 2020; Rao et al. 2025) (Fig. S1h) and contain the bacterial sterol mimics hopanoids alongside the eukaryotic sterols (Rao et al. 2025). So, what are the membrane features sensed by Mga2 and how has Mga2 evolved?

Our data show that both *S. pombe* and *S. japonicus* Mga2 proteins appear to sense lipid packing through the same rotational mechanism as the budding yeast homolog (Fig. 2d-f and Fig. S2a-d), but their activation thresholds have diverged (Fig. 2a-c). The species-specific activation patterns are encoded within the 55 amino acid region within the juxtamembrane domain of Mga2 (Fig. 2 and Fig. 3). The juxtamembrane domain, which includes the putative E3 ubiquitin ligase binding site and an intrinsically disordered region that extends to the TMH, is thought to act as a flexible linker that transmits the signal sensed by the TMH to the sites of ubiquitylation, promoting Mga2 processing (Ballweg et al. 2020). *S. pombe* and *S. japonicus* have three homologs of Rsp5 E3 ligase that polyubiquitylates Mga2 in budding yeast (Shcherbik et al. 2003; Covino et al. 2016; Ballweg et al. 2020). Suggesting that the ubiquitylation machinery and its target recognition are conserved, the deletion of the putative E3 ubiquitin ligase binding site abolishes Mga2 cleavage in both species and the *S. japonicus* Mga2 transplanted into *S. pombe* is capable of responding to oleic acid. We hypothesize that rather than differences in Mga2 ubiquitylation, it is the structural divergence in the juxtamembrane region, which alters the exposure of ubiquitinylated residues, setting *S. pombe* Mga2 more prone to cleavage than its *S. japonicus* counterpart.

Intrinsically disordered regions often evolve rapidly due to weaker structural constraints, thus serving as hotspots for rapid adaptation to changing conditions (Stormo et al. 2024). Of interest, the intrinsically disordered sequence within the identified 55 amino acid region is the most divergent part of the Mga2 C-terminus between all *Schizosaccharomyces* species, yet it is highly conserved within *S. pombe* natural isolates, suggesting that it remains functionally constrained in an individual species (Fig. 3e, f). Although these regions exhibit comparable degree of disorder, other sequence properties such as hydrophobicity (Park et al. 2025) may underlie the distinct *S. pombe* and *S. japonicus* Mga2 activation patterns. Tweaking an intrinsically disordered linker may allow the organisms to set different activation thresholds while maintaining the conserved mechanism of sensing.

Our results on lipidome adjustment during steady-state growth of both fission yeasts at different temperatures are consistent with glycerophospholipid FA asymmetry acting as an alternative to FA unsaturation to maintain membrane fluidity (Meyer and Bloch 1963; Panconi et al. 2023; Rao et al. 2025) (Fig. 1a, b). Of note, the asymmetrical glycerophospholipids assemble considerably thinner membranes in vitro (Rao et al. 2025). In line with that, a subset of *S. japonicus* single-pass transmembrane proteins functioning at the interface between the ER and other cellular organelles exhibit shorter TMHs as compared to other fission yeasts (Makarova et al. 2020). Despite the abundance of asymmetrical glycerophospholipids in *S. japonicus*, which do confer membrane disorder, Mga2 activation in this organism still depends on the degree of lipidome unsaturation (Fig. 2a, b). We favour a hypothesis that asymmetrical saturated and symmetrical unsaturated glycerophospholipids may be enriched in different nanodomains within the ER membrane, and the relatively long TMH of Mga2 (22 amino acids) may partition it into the thicker nanodomains containing symmetrical unsaturated glycerophospholipids to minimize hydrophobic mismatch. In this way, Mga2 would be sensitive specifically to its local environment. Additionally, asymmetrical saturated and symmetrical unsaturated glycerophospholipids may interact differently with hopanoids and ergosterol, contributing to the definition of membrane nanodomains with different physicochemical properties.

The highly abundant Ole1 in *S. pombe* – the expression of which is not controlled tightly by Mga2 – pushes the lipidome towards unsaturation. The high basal expression of *ole1* in this organism depends on the upstream non-coding region. Although *S. pombe* and *S. japonicus ole1* promoters share a similar organization of cis-regulatory elements (Cbf11 binding site-TATA-TSS), in *S. pombe* these elements lie within a low complexity region rich in poly(dA:dT) tracts that is absent from *S. japonicus* or other fission yeasts (Fig. 4c, Fig. S4c). Importantly, the *S. japonicus* promoter region fails to sustain Mga2-independent *ole1* expression at the levels required for *S. pombe* viability and growth at low temperatures (Fig. 4d-g and Methods). Genomic regions containing homopolymeric poly(dA:dT) tracts are prone to higher mutation frequencies (Tran et al. 1997; Ma et al. 2012). It is possible that slippage-mediated mutations in *S. pombe ole1* non-coding region have rearranged the sequence, increasing the strength of the core promoter and/or promoting chromatin accessibility. The larger 18kb locus upstream of *ole1* contains several *S. pombe*-specific genes (Fig. S4c), further suggesting that this genomic region constitutes a hotspot for evolutionary innovation.

We suggest that the high basal *ole1* expression that may have arisen due to local genome rearrangements during the evolution of *S. pombe*, has forced Mga2 to operate near the upper limit of the dynamic range of lipidome unsaturation. In this scenario, changes within the juxtamembrane region making Mga2 more prone to cleavage have allowed *S. pombe* to support high but not extreme levels of unsaturation; the latter incompatible with cell physiology (Fig. 4h).

When growing at different temperatures, *S. pombe* regulates the glycerophospholipid FA tail asymmetry – typically a minor feature of its lipidome – rather than regulating its abundant unsaturation. Conversely, *S. japonicus*, rich in asymmetrical glycerophospholipids, regulates its typically low FA unsaturation (Fig. 1a, b). Our work highlights an important design principle in homeostatic networks, which is the regulation of minor rather than major features subject to homeostasis. If the sensed cellular property is too abundant, the dynamic range of sensing collapses, with the system now functioning to prevent the overload (Fig. 4h). By demonstrating how conserved homeostatic circuits can be remodelled in evolution, this study advances our understanding of the mechanisms that confer robustness and adaptability to cellular regulatory networks.

## Materials and Methods

### Fission yeast strains, growth conditions and drug treatments

The *S. pombe* and *S. japonicus* strains used in this work are listed in Supplemental Data 4, Table 1. All strains were prototrophic. Standard fission yeast media and culture methods were used (Moreno et al. 1991; Furuya and Niki 2009; Aoki et al. 2010; Petersen and Russell 2016). For liquid cultures, cells were routinely grown in rich Yeast Extract with Supplements medium (YES) or Edinburgh minimal medium (EMM) containing the following supplements: adenine (93.75 mg/l), uracil (75 mg/l), histidine (75 mg/l) and leucine (75 mg/l) in 200 rpm shaking incubators at 30°C, unless otherwise stated. Typically, cells were pre-cultured in YES or EMM over eight hours, followed by dilution to appropriate OD_595_ and sub-culture overnight to reach mid-exponential phase (OD_595_ 0.4-0.6) the following morning. For cell treatment with oleic acid (C18:1, Sigma-Aldrich) a 3.2M oleic acid stock was prepared in DMSO, which was used to prepare a diluted 250 mM oleic acid stock in YES. The resulting solution was incubated for at least 1 hour at room temperature on a rocking platform to allow complete mixing of the fatty acid in the media. Final oleic acid concentration added to the cultures was 0.5 mM, except for the experiments involving the use of *S. pombe* Mga2 sensor constructs (Fig. 3 and Fig. S3), which required 1 mM oleic acid final concentration due to the lack of Mga2 transcriptional activator domain. Cerulenin (Sigma-Aldrich) treatment was performed by adding the drug to the cell cultures at a final concentration of 10 µM from a 10 mM stock in DMSO. Bortezomib (Sigma-Aldrich) was added to the cell cultures at a final concentration of 125 µM from a 10 mM stock in DMSO. *S. pombe* and *S. japonicus* mating was induced on SPA solid medium containing supplements as above at 25°C. Spores were dissected and germinated on YES agar plates using a dissection microscope (MSM 400, Singer Instruments).

### Materials

D-glucose anhydrous (cat. #G/0450/60) was purchased from ThermoFisher. Bacteriological agar (cat. #LP0011B) was purchased from Oxoid. Sodium phosphate dibasic dihydrate (cat. #71643), potassium hydrogen phthalate (cat. #P1088), ammonium chloride (cat. #A9434), adenine hemisulfate salt (cat. #A9126), L-histidine (cat. #H8000), L-leucine (cat. #L8000), uracil (cat. #U0750), sodium sulfate (cat. #239913), DMSO (cat. #D8418), oleic acid (cat. #O1008), cerulenin (cat. #C2389), bortezomib (cat. # 5.04314), trichloroacetic acid (cat. #T6399), acetone (cat. #179124), methanol (cat. #34860) and ethanol (cat. #32221) were purchased from Sigma-Aldrich. NEBridge Golden Gate Assembly Kit (cat. #E1601S) and NEBuilder HiFi DNA Assembly Master Mix (cat. #E2621L) were purchased from New England Biolabs. NuPage 4-12% BT gels (cat. #NP0321BOX), NuPage MOPS SDS running buffer (cat. #NP0001), NuPage transfer buffer (cat. #NP0006-1) and NuPage LDS sample buffer (cat. #NP0007) were purchased from Invitrogen. Nitrocellulose membranes 0.2 µm (cat. #1620112) were purchased from Bio-Rad. Mouse anti-GFP monoclonal antibody (cat. #11814460001) was purchased from Roche. Mouse anti-beta Actin monoclonal antibody (cat. #8224) was purchased from Abcam. Revert 700 Total Protein Stain Kit (cat. #926-11010) and IRDye 800CW Goat anti-Mouse IgG (cat. #926-32210) were purchased from LI-COR. RNeasy Plus Mini Kit (cat. #74134) and RNase-Free DNase Set (cat. #79256) were purchased from QIAGEN. Revertaid first strand cDNA synthesis kit (cat. #K1622) was purchased from Thermo Fisher. Probe Blue Mix Lo-ROX (cat. #PB20.21-01) was purchased from qPCRBIO.

### Molecular genetics

All primers are listed in Supplemental Data 4, Table 2. Molecular genetics manipulations were performed using PCR (Bähler et al. 1998)- or plasmid (Keeney and Boeke 1994)-based homologous recombination. A PCR-based method was used to knock out *S. pombe* and *S. japonicus mga2*, as well as to tag Ole1 at the C-terminus using *kanR* or *natR* as selection markers. Plasmids used to N-terminally tag *S. pombe* and *S. japonicus* Mga2 with GFP were obtained by Golden Gate cloning using NEBridge Golden Gate Assembly Kit (New England Biolabs). The rest of the plasmids used to obtain *S. pombe* and *S. japonicus* mutants were assembled using the Gibson Assembly Master Mix (New England Biolabs). All plasmids are listed in Supplemental Data 4, Table 3. All constructs were verified by sequencing. *S. japonicus* cells were transformed by electroporation (Aoki et al. 2010). *S. pombe* transformation was performed using lithium acetate and heat shock (Moreno et al. 1991). Transformants were selected on YES agar plates containing G418 (Sigma Aldrich) or nourseothricin (HKI Jena).

To test the viability of *S. pombe* mutants lacking a 1.6 kb or 2.8 kb fragment of the low-complexity region upstream of *ole1* ORF, a genomic copy of the 1.6 kb or 2.8 kb fragment was knocked out from *S. pombe* diploid cells that were obtained by heteroallelic complementation between the *ade6-M210* and *ade6-M216* mutant alleles as described in (Ekwall and Thon 2017). Diploid transformants were selected on Pombe Glutamate Medium (PMG) minus adenine containing twice the usual concentration of G418 (Sigma Aldrich). Viability of the resultant mutants was assayed by tetrad dissection (20 tetrads per deletion) and spore germination on YES agar plates, followed by replica-plating of the germinated spores on YES agar plates containing G418 (Sigma Aldrich). We found that 62/80 progeny were able to form colonies in the case of the mutant lacking 1.6 kb, from which 30/62 germinated spores were knockout mutants. Only 32/80 spores formed colonies in the case of the mutant lacking 2.8 kb, from which we recovered 0 knockout mutants.

### ESI-MS-based lipidomic analysis

Lipid standards were purchased from Avanti Polar Lipids (Alabaster, AL, USA). Solvents for extraction and MS analyses were liquid chromatography grade (Merck, Darmstadt, Germany) and Optima LC-MS grade (Thermo Fisher Scientific, Waltham, MA, USA), as applicable. All other chemicals were the best available grade purchased from Sigma-Aldrich or Thermo Fisher Scientific.

Exponentially growing *S. pombe* and *S. japonicus* cell cultures were grown in EMM at 30°C or the indicated temperature to mid-exponential phase and 10 ODs were harvested via filtration, then snap-frozen in liquid nitrogen. Cellular lipids were extracted and analysed using previously described methods (Makarova et al. 2020). Lipidomics data are expressed as mol% of polar lipids unless otherwise stated; polar lipids include all measured lipids except DG and TG. Double bond index (DBI), average chain length and average lipid species profile was calculated for the sum of major GPLs (PC, PI, PE and PS). DBI was calculated as Σ(db x [GPLi])/Σ[GPLi], where db is the total number of double bonds in fatty acyls in a given GPL species, and the square bracket indicates mol% of GPLs. Average chain length was calculated as Σ(C x [GPLi])/Σ[GPLi], where C is the total number of carbons in fatty acyls in a given GPL, and the square bracket indicates mol% of GPLs. ESI-MS quantification details for internal standards are provided in Supplemental Data 1, Table 4.

### RNA isolation

*S. pombe* and *S. japonicus* cell cultures were grown in EMM unless otherwise stated to mid-exponential phase and 10 ODs were collected. Total cellular RNA was extracted using RNeasy Plus Mini Kit (QIAGEN). Removal of genomic DNA from samples was performed during purification by on-column DNA digestion using RNase-Free DNase Set (QIAGEN).

### RNA sequencing and analysis

Strand-specific mRNA-seq libraries for the Illumina platform were generated and sequenced at the Genomics Science Technology Platform (STP) at the Francis Crick Institute. mRNA libraries were prepared using NEBNext Ultra II Directional PolyA mRNA enrichment with the subsequent quality control using Agilent Technology TapeStation system. A 200-cycle single read sequence run was performed on the NovaSeq 6000 Illumina instrument.

Raw sequence data was mapped to the *S. pombe* and *S. japonicus* genome using a Galaxy server pipeline (https://usegalaxy.org/). Quality assessment of the reads was performed using FastQC module. Adapter sequences and low-quality bases were removed with the Cutadapt module, applying a minimum quality score cutoff of 20 and a minimum read length of 20 bp. Cleaned reads were aligned to the *S. pombe* and *S. japonicus* reference genomes using the HISAT2 module with default parameters. Alignment quality was assessed using the MarkDuplicate, Gene Body Coverage, Samtools IdxStats and the Gene Body Coverage modules. The resulting BAM files were sorted and indexed for downstream analysis. Read counts per gene were quantified using featureCounts module, using the corresponding GTF annotation files. Differential gene expression analysis was performed using DESeq2 module. Differentially expressed genes (DEGs) were defined as those with an adjusted p-value (Benjamini–Hochberg correction) < 0.05 and an absolute log_2_ fold change ≥ 0.5 (Fig. 1c, d and Fig. 3b). Data were analysed using “DESeq2” R package (Love et al. 2014). Functional enrichment analyses were performed using g:Profiler (Kolberg et al. 2023). Note that functional enrichment analysis of *S. japonicus* DEGs was inferred using *S. pombe* Gene Ontology annotations via orthology mapping (69% of genes were mapped). Genes with fold change log_2_ values higher than 0.32 or lower than –0.32 and p values < 0.05 were considered for the analysis of commonly regulated genes (Fig. 1e). Data were plotted using “ggplot2” and “ComplexHeatmap” R packages (Gu et al. 2016; Wickham 2016; Gu 2022). Lists of DESeq2 results are presented in Supplemental Data 2, Tables 1, 2.

### Reverse transcription and real-time quantitative PCR (RT-qPCR)

Reverse transcription was performed as previously described in (Makarova et al. 2020) using oligo (dT) with First Strand cDNA Synthesis Kit (Roche). qPCRBIO Probe Blue Mix Lo-ROX was used for the real-time qPCR (RT-qPCR) assay with primers generated using Primer-BLAST tool (Ye et al. 2012) from the National Institute of Health (NIH). RT-qPCR was performed on a LightCycler 96 Instrument (Roche Diagnostics) in three biological and three technical repeats. For relative quantifications, RT-qPCR signal was normalized to actin expression levels. For absolute quantification of *sfGFP* mRNA abundance, a standard curve was generated using 1:10 serial dilutions of the digested plasmid pSO895, ranging from 10^8^ to 10^2^ ng/µl.

### Western blotting

*S. pombe* and *S. japonicus* cell cultures were grown in YES to mid-exponential phase and 5 ODs were pelleted for 1 min at 2103 x g and liquid medium was removed. Cells were resuspended in 1 ml ice-cold dH_2_O and transferred to 1.5 ml Eppendorf tubes. Cells were washed in 1 ml water and cell pellets were snap frozen in liquid nitrogen. Next, 110 µl ice-cold TCA was added and proteins were precipitated for 1 hour on ice. Lysates were pelleted for 10 minutes at 18213 x g at 4°C in an Eppendorf centrifuge. Pellets were washed once with 1ml ice-cold acetone. After removing the supernatant, pellets were dried in a speed-vac (Eppendorf) for 2 minutes at room temperature. Pellets were resuspended in 300 µl boiling buffer (50 mM Tris pH 8.0, 1 mM EDTA, 1% SDS) and transferred to pre-chilled screw-cap tubes with zirconium beads. Lysates were disrupted using a MP Biomedicals cell disruptor for 2 x 15 sec at 4°C and 6.5 m/sec with 2 minutes of cooling down on ice in between. Lysates were extracted from the beads by piercing the tubes using a hot needle and spun down for 3 min at 526 x g at 4°C. Debris were removed by spinning down lysates at 1383 x g for 4 minutes at room temperature. Lysates were then boiled at 95°C for 5 minutes. Next, 100 µl of 4x sample buffer with 10% β-mercaptoethanol was added and lysates were heated at 65°C for 10 minutes. 20 µl of lysate was loaded per lane on a NU-PAGE 4-12% Bis-Tris gel (Invitrogen) and run at 130V for 2.5 hours. Proteins were transferred to 0.2 µm membranes (Bio-Rad) at 100V for 1 hour. After transfer, membranes were blocked for 1 hour with TBST buffer containing 5% skimmed milk and then incubated overnight with mouse α-GFP (Roche) or mouse α-beta Actin (Abcam) antibodies. Membranes were washed with TBST and incubated for 1 hour in TBST buffer with IRDye800 conjugated α-mouse antibody (LI-COR Biosciences). Total protein levels were detected using the LI-COR Revert 700 Total Protein stain kit (LI-COR Biosciences). Proteins were detected using a Chemidoc MP System (Bio-Rad). Samples were collected as at least three biological replicates. Quantification was performed using Fiji (Schindelin et al. 2012).

### Cell growth assays

For growth rate determination *S. pombe* and *S. japonicus* cells were grown overnight in YES at 30°C or the indicated temperature until early-exponential phase. For experiments in the presence of oleic acid (Sigma-Aldrich), treatment was performed as indicated above. Cultures were then diluted to OD_595_ 0.1-0.15 with the same medium and loaded into a 96-well plate. Growth was measured every 60 minutes at the corresponding temperature using VICTOR Nivo multimode plate reader (PerkinElmer). Growth rates were calculated using the Growthcurver R package (Sprouffske and Wagner 2016). Experiments were repeated at least three times.

### Microscopy and image analysis

Prior to imaging, 1ml cell culture was concentrated to 50 μl by centrifugation at 1500 x g for 1 min. 2 μl cell suspension was loaded under a 22 × 22 mm glass coverslip (VWR, thickness: 1.5). Fluorescence images were acquired using Yokogawa CSU-X1 spinning disk confocal system with Eclipse Ti-E Inverted microscope with Nikon CFI Plan Apo Lambda 100× Oil N.A.=1.45 oil objective, 600 series SS 488nm SS 561nm lasers and Andor iXon Ultra U3-888-BV monochrome EMCCD camera controlled by Andor IQ3. Single plane images with inverted LUT (look-up-table) are shown.

Image analysis was performed using Fiji (Schindelin et al. 2012). Within the same experiment and for the same species, images are directly comparable as they are adjusted to equal brightness and contrast levels.

### Genetic crosses between *pole1^S.p.^*/*pole1^S.j.^* and *mga2Δ S. pombe* mutants

We dissected 27 tetrads (108 spores) from genetic crosses between *S. pombe pole1^S.p.^* (SO9614) or *pole1^S.j.^* (SO9573) and *mga2Δ* (SO8632) strains. If the double mutant from the corresponding cross were viable, the expected double mutant progeny would be 27/108 (25%). We found that crosses between the single mutants led to overall high spore lethality, likely due to a combination of lethality of the double mutants, or sporulation defects of the single mutants. In crosses between *pole1^S.p.^* and *mga2Δ*, 65/108 progeny were able to form colonies and out of 65 germinated spores, we recovered 12 double mutants. In crosses between *pole1^S.j.^*and *mga2Δ* mutants 44/108 spores germinated, from which we recovered 0 double mutants.

### Serial dilution growth assays

*S. pombe* cells were pre-cultured overnight in YES at 30°C until early-exponential phase. Cultures were then diluted to OD_595_ 0.1, and serial 10-fold dilutions were spotted on YES agar plates. Plates were incubated at the indicated temperatures for three days (24°C, 32°C and 36°C) or five days (16°C). Plates were then scanned using a Chemidoc MP System (Bio-Rad). Experiment was repeated three times with comparable results, and a representative experiment is shown in Fig. 4g.

### Bioinformatics

*S. pombe* and *S. japonicus* genomic and protein sequences were downloaded from Pombase (Rutherford et al. 2024) and JaponicusDB (Rutherford et al. 2022), respectively. Sequences from other yeasts were downloaded from Ensembl Fungi (Andrew et al. 2022). We noticed that *S. japonicus* Mga2 protein sequence was truncated in JaponicusDB and Ensembl Fungi because of a wrongly predicted intron. To obtain the correct sequence, we performed random-primer reverse transcription followed by PCR amplification and sequencing of the resulting cDNA amplicon. We did not detect any evidence for splicing. It was confirmed by the analysis of RNA-seq read coverage. The corrected protein sequence can be found in Supplemental Data 4, Table 4. Enrichment analysis of putative Cbf11-binding motifs was performed using Simple Enrichment Analysis (SEA) tool from MEME suite (Bailey and Grant 2021) with default parameters. Sequences were aligned using Clustal Omega Multiple Sequence Alignment tool (Madeira et al. 2024) using default parameters and represented using ESPript 3.0 (Robert and Gouet 2014). Transmembrane helices were predicted using TMHMM 2.0 (Krogh et al. 2001) using default parameters. Natural variation in *S. pombe* for *mga2* were retrieved from a SnpEff-annotated VCF file of 161 *S. pombe* isolates (including 57 non-clonal strains) from (Jeffares et al. 2015). The VCF file was downloaded from figshare (https://figshare.com/articles/dataset/SNP_calls_for_all_161_strains_/3978303?file=6217275). BCFtools (Danecek et al. 2021) (version 1.11) index was used to create an index file for the VCF file. BCFtools query was used to extract genetic variation for *mga2* and predicted effects from the following genomic range: chr1:4125212-4128667. The bash scripts to download, compress and create index file (prepareinputs.sh) and to retrieve variation (variantexplorer.sh) are available on the following git repository: https://github.com/billardb/pombe-variant-explorer.git. Single nucleotide variants located in *mga2*, along with their predicted effects are presented in Supplemental Data 3, Table 1.

### Statistics and reproducibility

The statistical details of experiments, including the number of biological and technical replicates and the dispersion and precision measures can be found in Figure Legends, Supplementary Figure Legends and Materials and Methods. All data were analysed using unpaired t-test statistical analysis, unless indicated otherwise. All plots were generated using GraphPad Prism 10. No data were excluded in the cell biological and physiological experiments.

## Competing interests

The authors declare no competing or financial interests.

## Acknowledgements

We are grateful to the Oliferenko lab for discussions and Damien Coudreuse and Eugene Makeyev for advice and suggestions on the manuscript. Many thanks to Elizabeth Carter for media preparations and the Crick Genomics STP for RNA-seq experiments. E. G. G. has been supported through a long-term EMBO postdoctoral fellowship (ALTF 712-2022, E. G. G.) and the UKRI guarantee of a MSCA postdoctoral fellowship (EP/Y024702/1, E. G. G). B. B. was supported by a grant from the Agence Nationale de la Recherche (PRC eVOLve, ANR-18-CD13-0009) to Damien Coudreuse. We thank the Single Cell Omics Advanced Core Facility staff of the HCEMM and Biological Research Center for help with their resources and their support. HCEMM has received funding from the EU’s Horizon 2020 research and innovation program under grant agreement No. 739593 and KIM NKFIA 2022-2.1.1-NL-2022-00005. This work was supported by the Francis Crick Institute, which receives its core funding from Cancer Research UK (CC0102), the UK Medical Research Council (CC0102), and the Wellcome Trust (CC0102), and the Wellcome Trust Senior Investigator Award (103741/Z/14/Z) and Wellcome Trust Investigator Award in Science (220790/Z/20/Z) to S. O.

## Author contributions

E. G. G. conceived and performed cell biological, biochemical experiments and bioinformatics analyses; generated strains; analyzed data; and co-wrote the manuscript. P. G. conceived and performed cell biological and gene expression experiments; generated strains; analyzed data; and edited the manuscript. Y. G. determined the correct sequence for *S. japonicus* Mga2; generated strains; and edited the manuscript. S. F. performed initial characterisation of GFP-Mga2 and *mga2Δ* strains; and edited the manuscript. B. B. wrote a script to retrieve *S. pombe* genomic variants; and edited the manuscript. G. B. and M. P. designed, performed, and interpreted ESI-MS lipidomics experiments and edited the manuscript. S. O. conceived and interpreted experiments and co-wrote the manuscript. This research was funded in whole, or in part, by the Wellcome Trust (103741/Z/14/Z; 220790/Z/20/Z to S. O.). For the purpose of Open Access, the author has applied a CC-BY public copyright licence to any Author Accepted Manuscript version arising from this submission.

---

**Figure S1.**
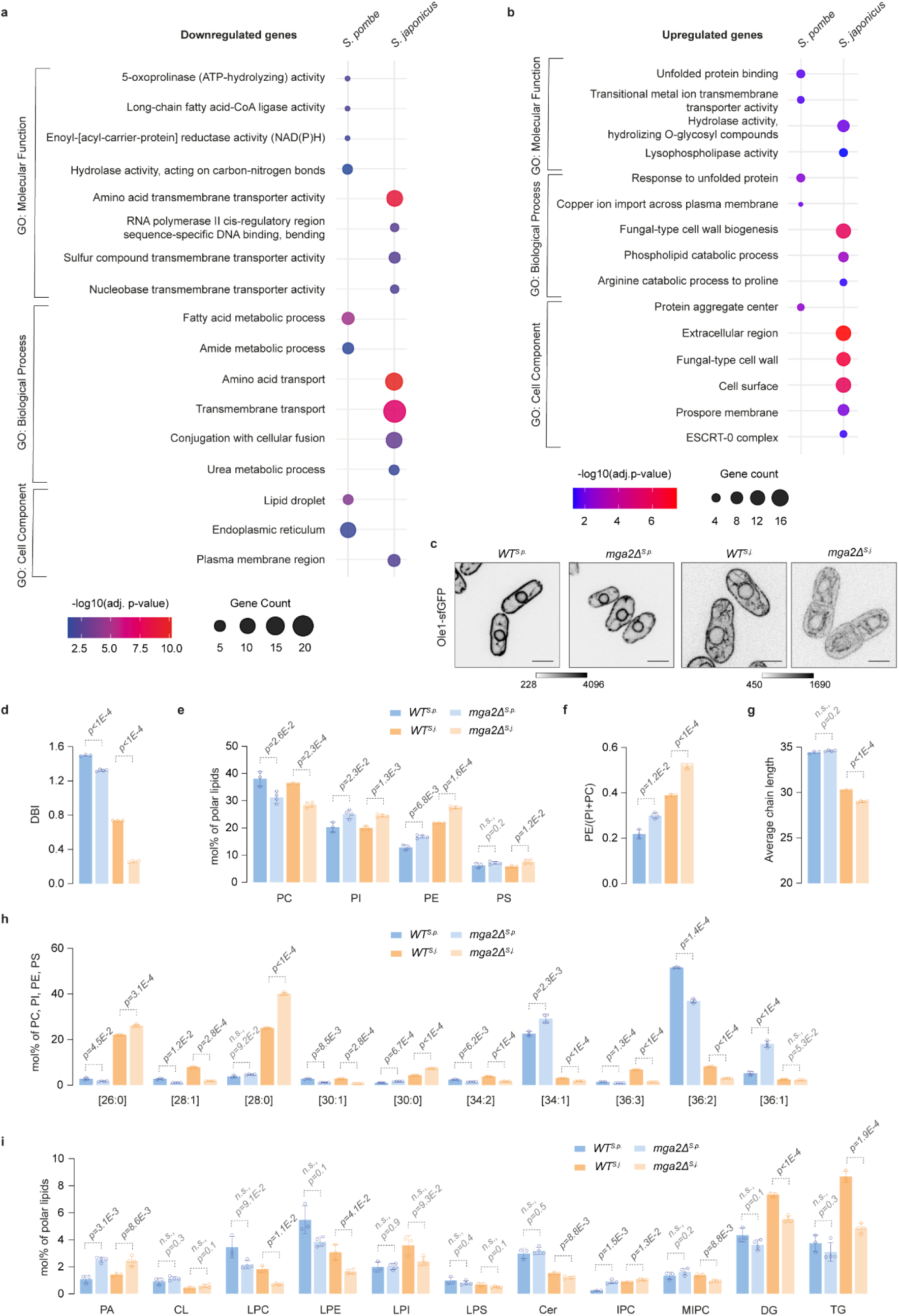
Related to Figure 1. The loss of Mga2 results in decreased fatty acid unsaturation and further remodelling of the cellular lipidome in both fission yeasts. (**a**, **b**) Gene Ontology (GO) enrichment analyses of differentially downregulated (**a**) and upregulated (**b**) genes in *mga2Δ* mutant vs wild type in *S. pombe* and *S. japonicus* cells grown in EMM (-0.5 > log_2_(fold change) > 0.5 and adjusted p-value < 0.05), performed using g:Profiler. The size of each dot represents the number of terms associated with the GO term, and the colour represents the adjusted p-value. (**c**) Single plane spinning disk confocal images of *S. pombe* and *S. japonicus* wild type (*WT*) and *mga2Δ* cells expressing Ole1-sfGFP grown in YES. Scale bars represent 5 µm. All images were contrasted to the same scale. Greyscale bar is included. (**d**) Comparison of double bond indices (DBI) calculated for the sum of the four main glycerophospholipid (GPL) classes ((PC), phosphatidylinositol (PI), phosphatidylethanolamine (PE) and phosphatidylserine (PS)) in *S. pombe* and *S. japonicus WT* and *mga2Δ* strains grown in EMM. (**e**) Relative abundance of the four main GPL classes in *S. pombe* and *S. japonicus WT* and *mag2Δ* cells grown in EMM. (**f**) Ratio between PE and the sum of PC and PI (PE/(PC+PI)) calculated from the ESI-MS lipidomics data shown in (**e**). (**g**) Average combined FA length calculated for the sum of PC, PI, PE, and PS in *S. pombe* and *S. japonicus WT* and *mga2Δ* strains grown in EMM. (**h**) Molecular species composition calculated for the sum of PC, PI, PE, and PS in *S. pombe* and *S. japonicus WT* and *mga2Δ* cells grown in EMM. The categories are shown as the total number of carbon atoms: total number of double bonds in acyl chains. (**i**) Relative abundance of the indicated lipid classes: phosphatidic acid (PA), cardiolipin (CL), lysophosphatidylcholine (LPC), lysophosphatidylethanolamine (LPE), lysophosphatidylinositol (LPI), lysophosphatidylserine (LPS), ceramide (Cer), inositol phosphoceramide (IPC), mannosyl-inositolphosphoceramide (MIPC), diacylglycerol (DG) and triacylglycerol (TG) in *S. pombe* and *S. japonicus WT* and *mga2Δ* cells grown in EMM. (**d**-**i**) Data are represented as average ± S.D. (n=2 biological, 1 technical repeats for *WT*; 2 biological, 2 technical repeats for *mga2Δ*). p-values are derived from two-tailed unpaired t-test.

**Figure S2.**
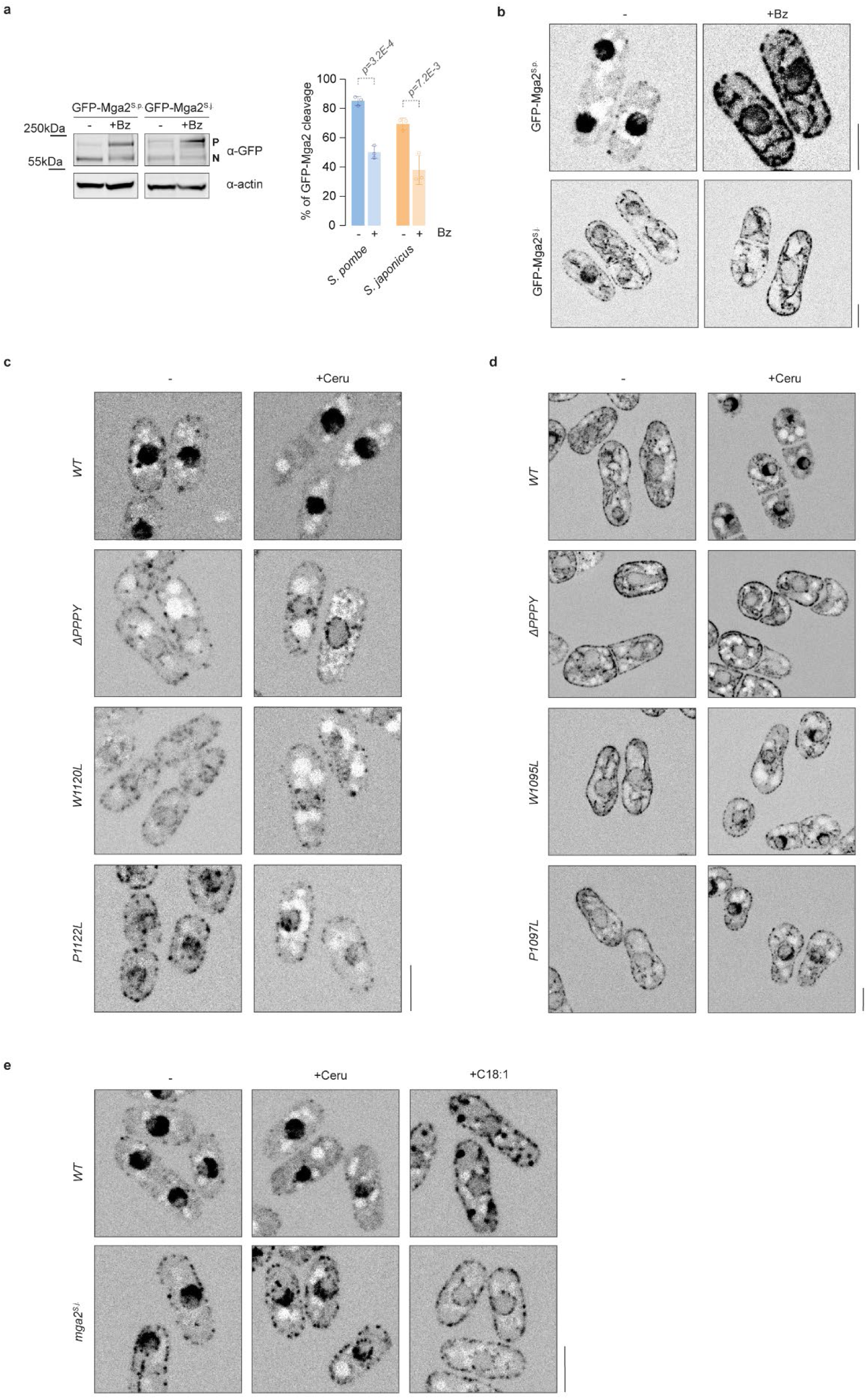
Related to Figure 2. Mga2 is proteolytically activated via the proteasome in both sister species. (**a**) Western blot of *S. pombe* and *S. japonicus* cells expressing GFP-Mga2 grown in YES in the absence or the presence of 125 µM of the proteasomal inhibitor Bortezomib (Bz) for 3.5 hours. Actin was used as a loading control. P and N indicate precursor and cleaved N-terminal transcriptional activator forms, respectively. The right panel shows the quantification of GFP-Mga2 cleavage, calculated as N / (N + P), expressed as percentage. Data are represented as average ± S.D. (n=3 biological repeats). p-values are derived from two-tailed unpaired t-test. (**b**) Single plane spinning disk confocal images of *S. pombe* and *S. japonicus* cells from experiment shown in (a). (**c**, **d**) Single plane spinning disk confocal images of *S. pombe* (**c**) and *S. japonicus* (**d**) cells from experiments shown in Fig. 2e and 2f, respectively. (**e**) Single plane spinning disk confocal images of *S. pombe* cells from experiment shown in Fig. 2g. (**b**-**e**) Scale bars (5 µm) for each species shown bottom right.

**Figure S3.**
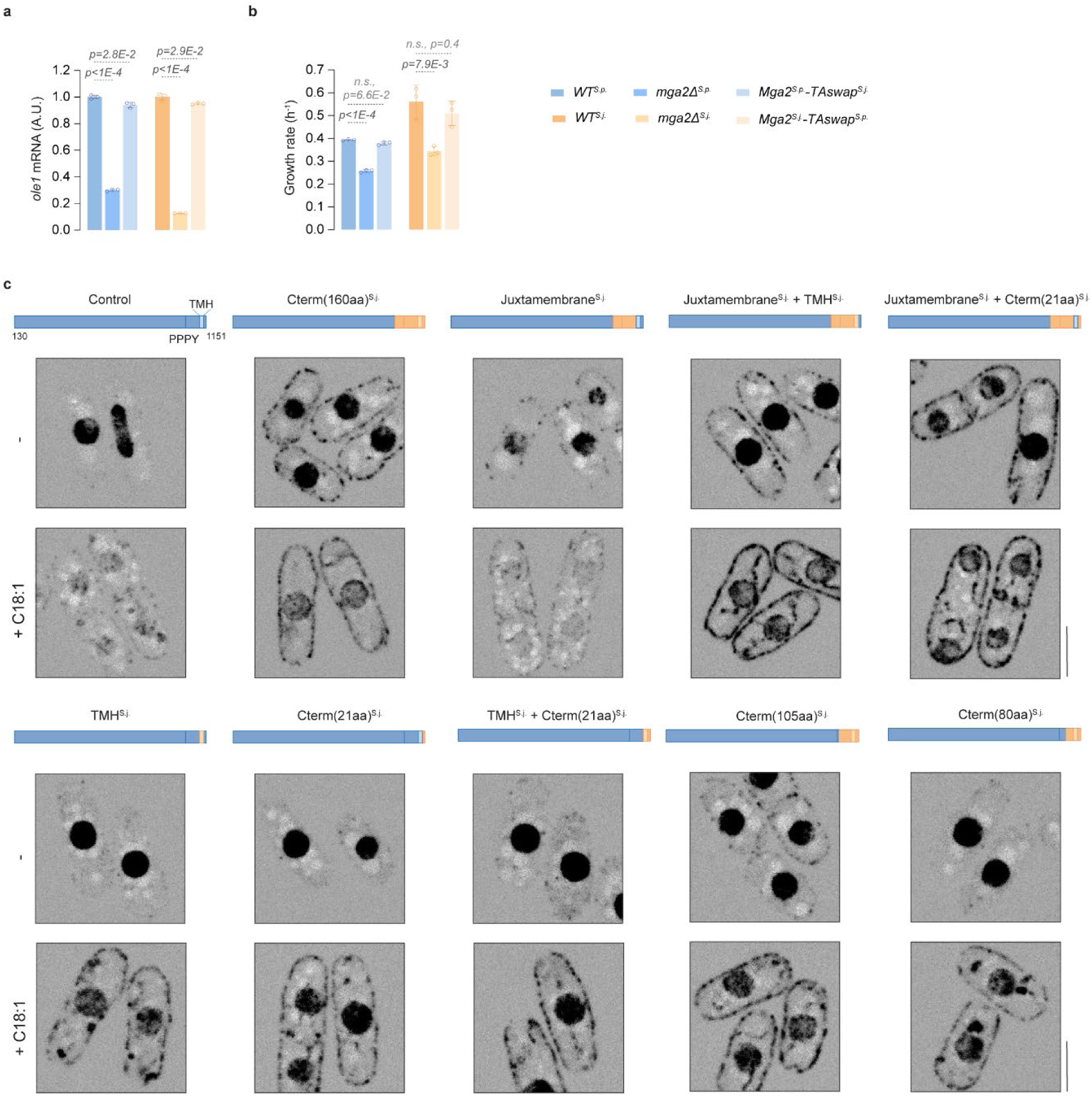
Related to Figure 3. Mga2 transcriptional activator domain is functionally interchangeable between *S. pombe* and *S. japonicus*. (**a**) Steady-state *ole1* mRNA levels in *S. pombe* and *S. japonicus* strains of the indicated genotypes grown in YES, as measured by qPCR, normalized to the wild type (*WT*). (**b**) Comparison of growth rates of *S. pombe* and *S. japonicus* strains of the indicated genotypes grown in YES. (**c**) Single plane spinning disk confocal images of *S. pombe* cells from the experiment in Fig. 3c, d. Schematics of the corresponding constructs are shown above each image. Scale bars represent 5 µm. (**a**, **b**) Data are represented as average ± S.D. (n=3 biological repeats). p-values are derived from two-tailed unpaired t-test.

**Figure S4.**
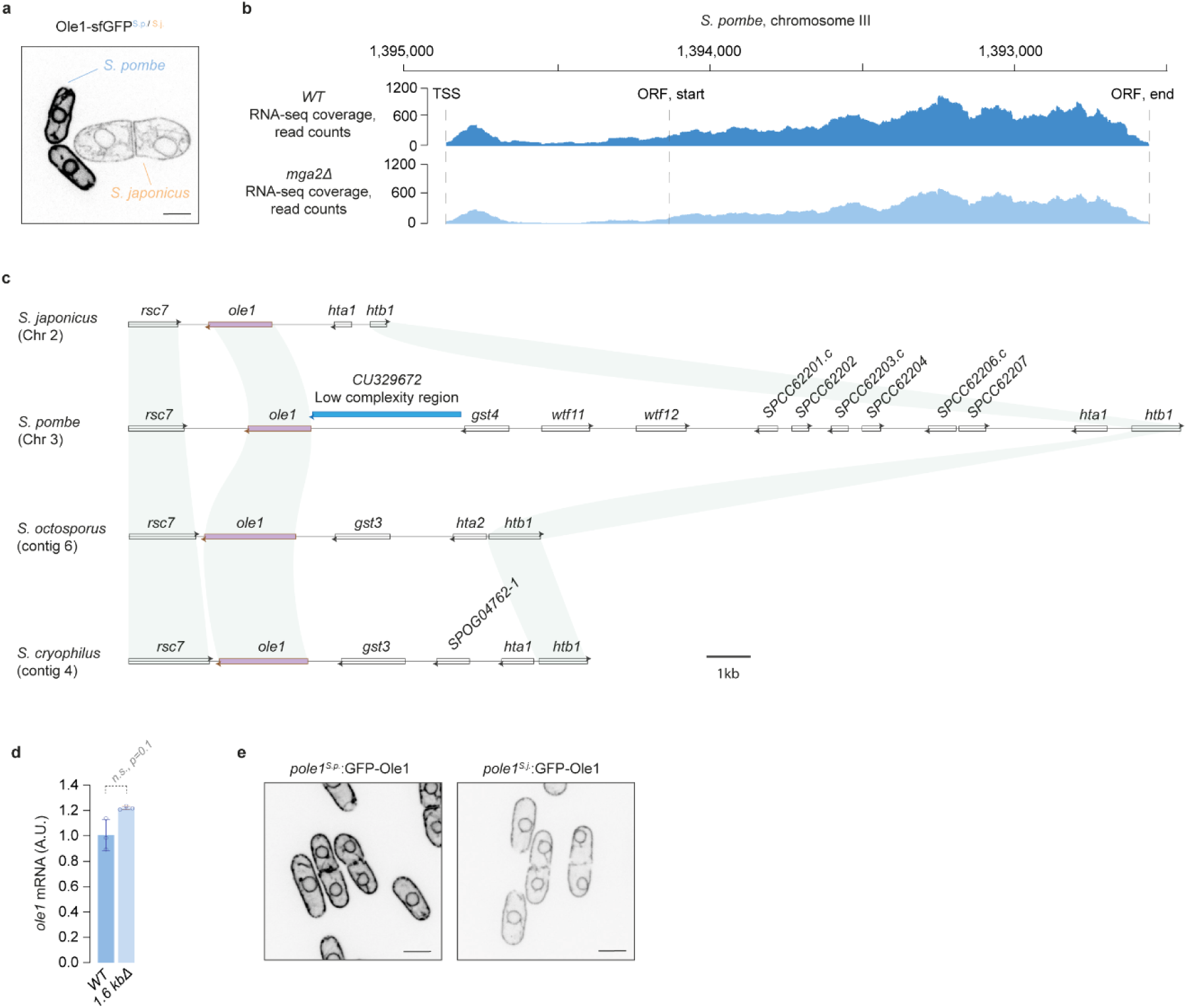
Related to Figure 4. *S. pombe ole1* intergenic region shows major genetic rearrangement. (**a**) A single plane spinning disk confocal image of *S. pombe* and *S. japonicus* cells expressing Ole1-sfGFP grown in YES. (**b**) Mapping of *ole1* transcript in *S. pombe* wild type and *mga2Δ* cells, IGV viewer is shown. (**c**) Diagram of the region between *rsc7* and *htb1* in *S. japonicus*, *S. pombe*, *S. octosporus* and *S. cryophilus*. The *ole1* gene is shown in purple. *S. pombe* low complexity region upstream of *ole1* is shown in blue. (**d**) Steady-state *ole1* mRNA levels in *S. pombe* wild type (*WT*) and the mutant lacking 1.6 kb of the low complexity region *Δ1.6 kb* grown in YES, as measured by qPCR, normalized to the *WT*. Data are represented as average ± S.D. (n=3 biological repeats). p-values are derived from two-tailed unpaired t-test. (**e**) Single plane spinning disk confocal images of *pole1^S.p.^*:GFP-Ole1 and *pole1^S.j.^:*GFP-Ole1 *S. pombe* cells, grown in YES. (**a**, **e**) Scale bars represent 5 µm.

